# Elevated N-glycosylation of immunoglobulin G variable regions in myasthenia gravis highlights a commonality across autoantibody-associated diseases

**DOI:** 10.1101/2021.03.06.434199

**Authors:** Caleigh Mandel-Brehm, Miriam L. Fichtner, Ruoyi Jiang, Valerie J. Winton, Sara E. Vazquez, Minh C. Pham, Kenneth B. Hoehn, Neil L. Kelleher, Richard J. Nowak, Steven H. Kleinstein, Michael R. Wilson, Joseph L. DeRisi, Kevin C. O’Connor

**Author notes:** Corresponding author: Kevin C. O’Connor, Ph.D. Departments of Neurology and Immunobiology Yale University School of Medicine. CMB, MLF and RJ contributed equally as first authors. JLD and KCO contributed equally as senior authors.

## Abstract

Elevated N-linked glycosylation of immunoglobulin G variable regions (IgG-V^N-Glyc^) is an emerging molecular phenotype associated with autoimmune disorders. To test the broader specificity of elevated IgG-V^N-Glyc^, we studied patients with distinct subtypes of myasthenia gravis (MG), a B cell-mediated autoimmune disease. Our experimental design included adaptive immune receptor repertoire sequencing to quantify and characterize N-glycosylation sites in the global B cell receptor repertoire, proteomics to examine glycosylation patterns of the circulating IgG, and production of human-derived recombinant autoantibodies, which were studied with mass spectrometry and antigen binding assays to confirm occupation of glycosylation sites and determine whether they alter binding. We found that the frequency of IgG-V^N-Glyc^ motifs was increased in the B cell repertoire of MG patients when compared to healthy donors. Motifs were introduced by both biased V gene segment usage and somatic hypermutation. IgG-V^N-Glyc^ could be observed in the circulating IgG in a subset of MG patients. Autoantigen binding, by patient-derived MG autoantigen-specific monoclonal antibodies with experimentally confirmed presence of IgG-V^N-Glyc^, was not altered by the glycosylation. Our findings extend prior work on patterns of variable region N-linked glycosylation in autoimmunity to MG subtypes. Although occupied IgG-V^N-Glyc^ motifs are found on MG autoantigen-specific monoclonal antibodies, they are not required for binding to the autoantigen in this disease.

## Introduction

The vast diversity of immunoglobulin G variable regions (IgG-V) is critical for host immunity. This diversity arises through VDJ recombination and somatic hypermutation (SHM). Historically, IgG-V diversity has been represented by amino acid sequence alone with little focus on post-translational modifications. Recently, the presence of N-linked glycosylation in IgG-V (IgG-V^N-Glyc^) has been shown to contribute to diversity (1, 2). IgG-V^N-Glyc^ is contingent upon the presence of the predictive N-Glyc amino acid motif N-X-S/T, where X can be any amino acid except for proline. This motif is most often introduced as a consequence of SHM (3). Less often it can be provided by the few germline gene segments (IGHV1-8, IGHV4-34, IGHV5-10-1, IGLV3-12, and IGLV5-37) in which it is encoded (4).

The percentage of IgG in healthy individuals that includes V-region glycosylation is approximately 15-25%; the range reflects different approaches of measurement (1). The occurrence of IgG-V^N-Glyc^ also varies among the IgG subclasses, with skewing toward higher frequencies in antibodies of the IgG4 subclass (5). Higher frequencies of IgG-V^N-Glyc^ than that which is found in healthy individuals have been observed in B cell malignancies (6–10) and in autoimmune diseases (11). Specifically, increased frequencies have been reported for ANCA-associated vasculitis (AAV) (12–14), rheumatoid arthritis (RA) (15–18), and primary Sjogren’s syndrome (pSS) (19, 20). The *in vivo* function of glycosylation in the IgG-V, a critical region of antigen contact, is not thoroughly understood. Follicular lymphomas may leverage N-glycosylation of their B cell receptors to activate antigen-independent signaling pathways that support survival (21). Antigen binding can also be influenced by IgG-V^N-Glyc^; this includes both increases and decreases in affinity and modulated functional activity. This is well highlighted by anti-citrullinated protein autoantibodies found in RA patients, where 80-100% of the autoantibodies include IgG-V^N-Glyc^, and binding is consequently altered (5, 16, 17).

Myasthenia gravis (MG) is an autoimmune disorder affecting neuromuscular transmission. MG patients experience severe muscle weakness and increased fatigability (22, 23). The molecular immunopathology of MG is directly attributed to the presence of circulating IgG isotype autoantibodies specifically targeting extracellular domains of postsynaptic membrane proteins at the neuromuscular junction (NMJ) (23, 24). The most common subtype of autoantibody-mediated MG (approximately 85% of patients) is characterized by autoantibodies against the nicotinic acetylcholine receptor (AChR) (23). In many of the remaining patients, autoantibodies targeting the muscle-specific kinase (MuSK) are present (25, 26). While both anti-AChR and anti-MuSK antibodies cause disease, the underlying immune pathophysiology of these two MG subtypes is distinct (27). AChR MG is governed primarily by IgG1 subclass autoantibodies that facilitate pathology through blocking acetylcholine, activating complement-mediated damage and initiating internalization of AChRs (28–31). Conversely, the MuSK MG subtype is most often associated with IgG4 subclass autoantibodies, which are incapable of activating complement, but rather mediate pathology through blocking MuSK binding partners and its kinase activity (32–34).

Given that IgG isotype autoantibodies directly facilitate MG pathology and that their divergent autoimmune mechanisms include different IgG subclasses (IgG1 and IgG4) known to include varying frequencies of IgG-V^N-Glyc^, we hypothesized that N-linked glycosylation might be differentially elevated in these two distinct MG subtypes. To that end we applied complementary sequencing and proteomic-based approaches to investigate IgG-V^N-Glyc^ patterns in AChR and MuSK MG. Nucleotide-level sequencing was used to test for elevated IgG-V^N-Glyc^ frequency in MuSK and AChR MG B cell receptor repertoires. Antibodies from sera were then evaluated with proteomic approaches to determine whether elevated IgG-V^N-Glyc^ could be observed in the circulation. Finally, we tested whether N-linked glycans impact binding to pathogenic targets by using patient-derived monoclonal autoantibodies with N-linked glycan occupancy validated by mass spectrometry. We show that IgG-V^N-Glyc^ are more frequent in both AChR and MuSK MG in comparison to healthy controls and that the patterns differ between the two MG subtypes. However, the presence of IgG-V^N-Glyc^ does not interrupt the binding of the pathogenic autoantibodies to their target antigens.

## Results

### The frequency of IgG-V^N-glyc^ is elevated in the B cell repertoire of patients with MG

N-linked glycosylation sites only occur at amino acid sequence positions with the motif (N-X-S/T, where X = not proline). Elevated IgG-V^N-Glyc^ in MG could arise from the introduction of these sites by SHM or the use of germline sites found in a small subset of VH gene segments (IGHV1-8, IGHV4-34 and IGHV5-10). To quantify global differences in the glycosylation frequency of the B cell repertoire, we examined the encoded B cell receptor repertoire generated by adaptive immune receptor repertoire sequencing (AIRR-seq) from the mRNA of circulating PBMCs from healthy donors (HD) and MG patients (**Table S1**). The MuSK MG patient cohort (N=3) included 12 unique timepoint samples, the AChR (N=10) included 10 unique timepoint samples, and each HD (N=9) included a single time point. The AIRR-seq library included a total of 10,565,778 (heavy chain only) raw reads; after quality control and processing, a high-fidelity data set was generated that consisted of 764,644 unique error-corrected sequences, which was further filtered to include only IgG subclass sequences that consisted of 232,094 sequences.

We observed a statistically significant elevation in median IgG-V^N-Glyc^ site frequency for AChR MG (13.0%; P=0.039, one-tailed Wilcoxon test) and MuSK MG (17.4%, P=0.018, one-tailed Wilcoxon test) in comparison to healthy controls (10.3%) (**Figure 1A**). To investigate if the increased frequency of N-linked glycosylation sites was generated through preferred use of the three VH-gene segments that encode an N-X-S/T motif or through SHM, we assessed the frequency of the motif in germline reversions of the VH-gene segments (**Figure 1B**). We observed no differences in the germline frequency of IgG-V^N-Glyc^ sites when comparing healthy and AChR MG patients (P=0.55, one-tailed Wilcoxon test), while the MuSK MG cohort exhibited a significant difference (P=0.05, one-tailed Wilcoxon test), thus reflecting increases in the usage of select V gene segments (IGHV1-8, IGHV4-34, IGHV5-10-1). An illustrative example of N-X-S/T motif acquisition and conservation through the SHM process is shown for a B cell clonal family present in a MuSK MG repertoire, which includes acquisition of two motifs (**Figure S1**).

**Figure 1.**
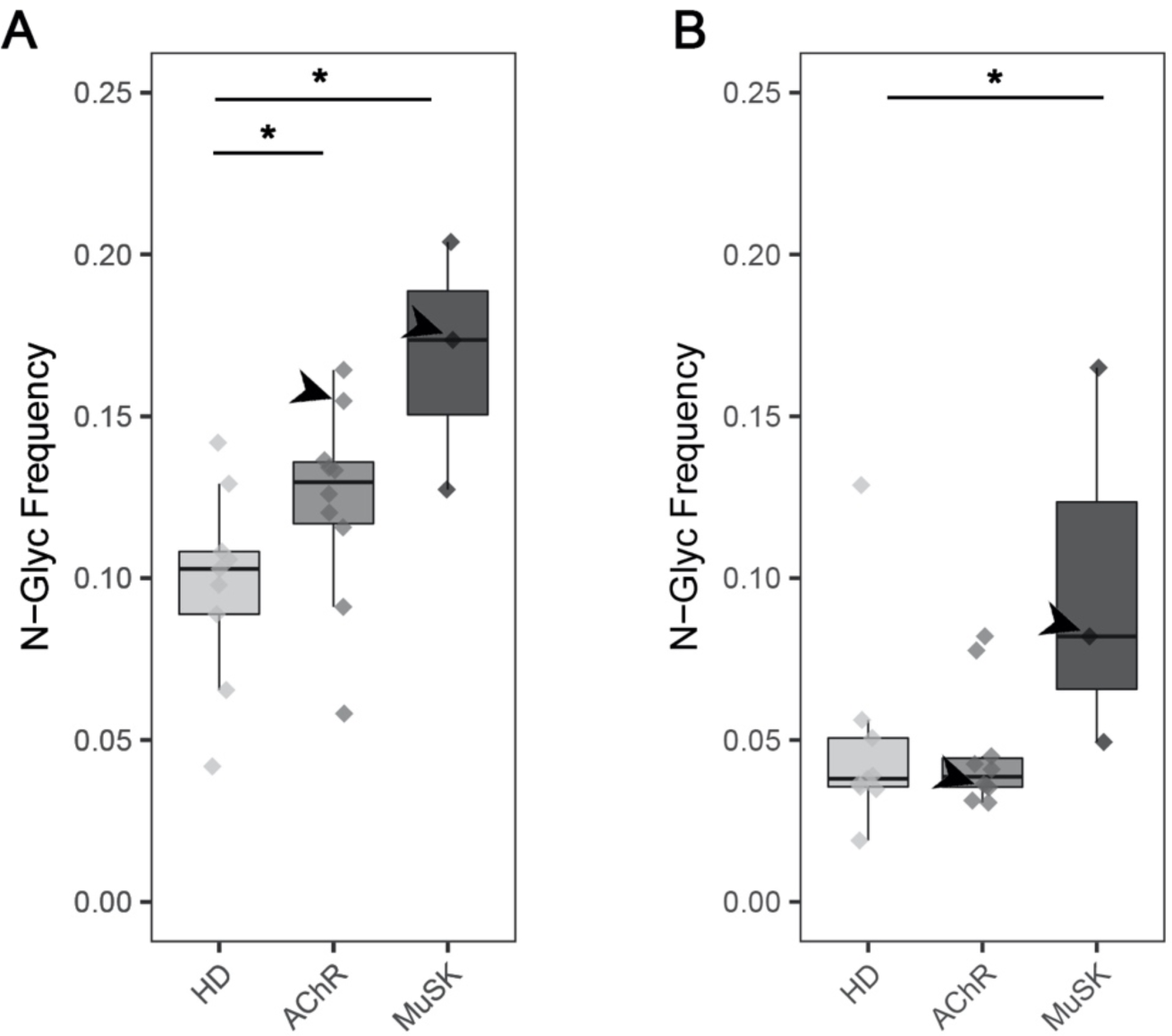
Frequency of IgG isotype-specific V^N-Glyc^ sites in the B cell receptor repertoire. Analysis of adaptive immune receptor repertoire sequencing showing the frequency of N-linked glycosylation motifs in AChR and MuSK MG BCR repertoires relative to HDs. (A) Frequency of N-linked glycosylation motifs (N-X-S/T) in the V gene sequence repertoire derived from healthy, AChR and MuSK MG patient repertoires is shown. (B) Frequency of N-linked glycosylation motifs (N-X-S/T, average number of sites per V(D)J sequence) in germline reverted V gene sequences from BCR repertoire derived from healthy, AChR and MuSK MG patient repertoires. A significance threshold of p <0.05 was used and shown on plots with a single asterisk. Arrows point to AChR MG-1 or MuSK MG-1 depending on the boxplot in (A) and (B).

We then examined if differences in glycosylation frequency were specific to complementarity-determining (CDR) regions, which are primarily responsible for antigen contact, or also included distribution in the framework regions (FWR), which maintain structural integrity of the variable domains. The motif could be found in all CDRs and FWRs in sequences from the HDs and MG patients with the exception of FWR2, in which the motif was absent in all sequences (**Figure S2A**). The motif was most often observed in the FWR3 sequences from the HDs and MG patients. Sequences from the AChR MG patients revealed a significant difference in motif frequency only in the CDR2 region in comparison to HD (P=0.047, one-tailed Wilcoxon-test). Elevated frequencies were observed in comparisons of MuSK MG and HDs at all regions (**Figure S2A**), and statistically significant differences were observed in the FWR1, CDR2, and FWR4 regions (P=0.009, P=0.0045, P=0.042, respectively - one-tailed Wilcoxon tests). Examining the location of the motifs within each region (**Figure S2B**) showed that they were present throughout, but were not uniformly distributed, as some areas showed enriched accumulation. Those present in FWR1 and FWR4, although rare, were found close to the CDRs that they neighbor, CDR1 and CDR3 respectively (**Figure S2B**).

In summary, the frequency of variable region N-linked glycosylation sites among IgG switched B cells differ when comparing healthy controls and AChR or MuSK MG patients. These differences result from SHM in AChR MG, while differences found in MuSK MG result from both SHM and elevated usage of V genes with germline encoded N-linked glycosylation sites.

### Proteomic analysis demonstrates elevated IgG-V^N-glyc^ in MG

Serum-derived IgG heavy chains associated with human autoimmunity, such as those in RA and ANCA-associated vasculitis, migrate at a higher molecular weight (MW) than those of healthy controls due to the presence of IgG-V^N-Glyc^ (14, 16). Having demonstrated that the B cell repertoire of both AChR and MuSK MG include elevated IgG-V^N-Glyc^ site frequency, we next sought to investigate if circulating IgG from patients with MG reflected this MW increase. To that end, we analyzed IgG purified from serum samples from the MG cohort (MuSK MG, N=3; AChR MG, N=9) for the presence of IgG-V^N-Glyc^ (**Table S1**). Longitudinal samples were also included to evaluate the temporal stability of IgG-V^N-Glyc^ patterns (**Table S1**). The IgG-V^N-Glyc^ presence was tested through the assessment of immunoglobulin heavy chain migration patterns by SDS-PAGE. Serum-derived IgG from a patient with RA was included as a positive control (**Figure 2A**). IgG migration patterns between healthy individuals and MG patients were compared (**Figure 2B**); differences were noted for one AChR patient (AChR MG-1) and one MuSK patient (MuSK MG-1). Longitudinal samples were assessed spanning a period of four years of clinical disease; the altered migration patterns remained consistent through all of the time points collected from these two subjects (**Figure 2C**). These two subjects also demonstrated an elevated frequency of IgG-V^N-glyc^ in their B cell repertoire (**Figure 1** **arrows)**.

**Figure 2.**
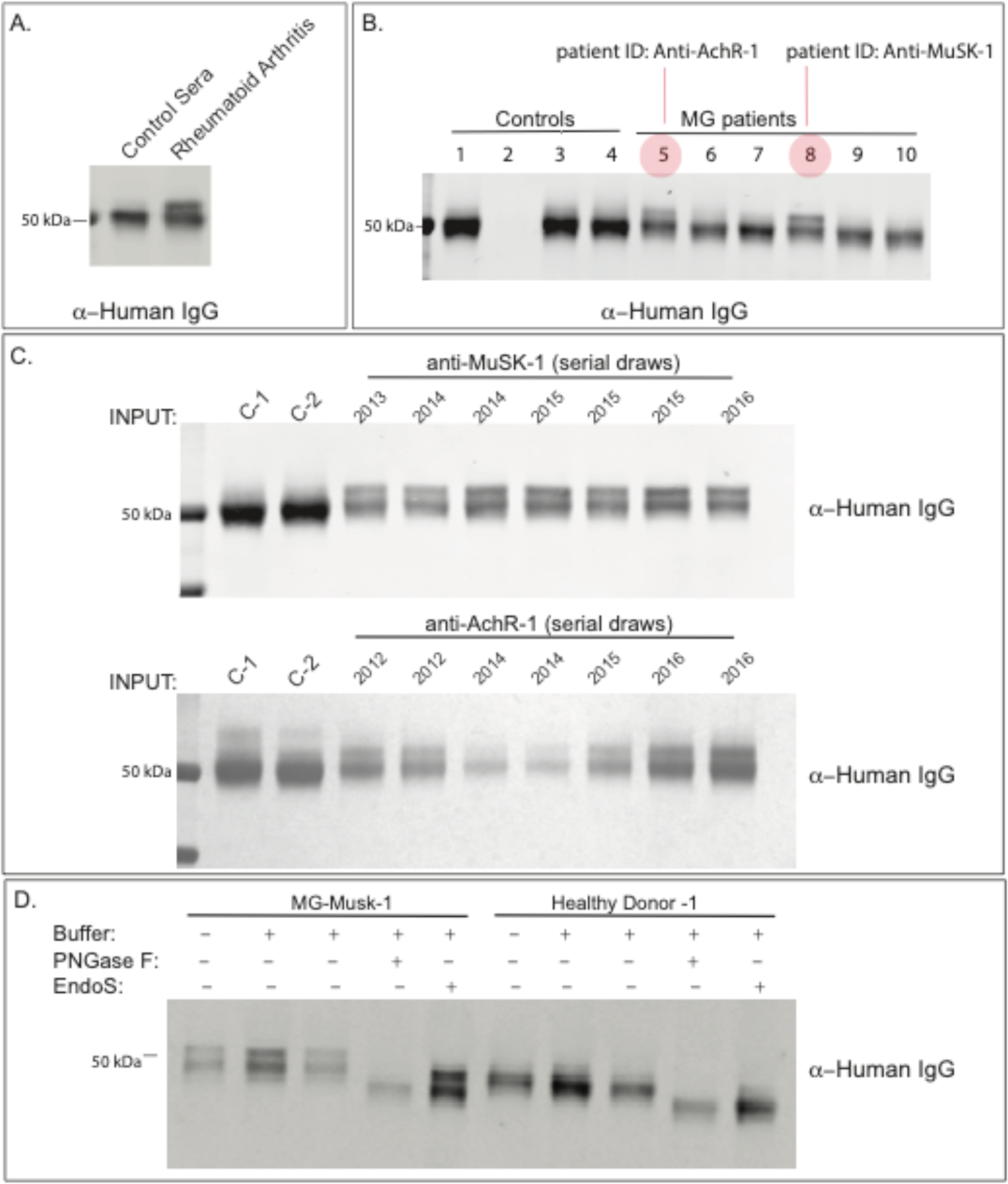
Proteomic analysis of glycosylation of serum IgG. Screen reveals evidence of elevated Fab N-linked glycosylation in myasthenia gravis. Serum IgG heavy chain migratory patterns from SDS-PAGE and immunoblotting with anti-human secondary are shown for sub-panels. (A) Serum IgG heavy chain migration pattern from non-inflammatory control and a patient with RA shown by immunoblot. (B) Serum IgG heavy chain migration patterns for non-inflammatory controls (Lane 1, 3, 4), A/G beads only (No antibody, Lane 2) and clinically confirmed AChR MG (Lanes 5, 6, 7) or MuSK MG (Lanes 8, 9, 10) shown by immunoblot. Lane 5 corresponds to subject AChR MG-1 and lane 8 corresponds to subject MuSK MG-1. (C) Longitudinal serum IgG heavy chain patterns from subjects MuSK MG-1 and AChR MG-1 shown by immunoblot. (D) Enzymatic validation of N-linked glycosylation shown by immunoblot. Schematic of N-glycan cleavage specificity by PNGase F and Endo S for IgG. Endo S only cleaves N-glycans at N297; PNGase F cannot cleave N-linked glycans at N297. Treatment with PNGase F but not Endo S results in loss of migration phenotype. RA = Rheumatoid Arthritis. Input = 10% of total immunoprecipitated antibody using A/G beads from 1 ul of patient sera.

To assess whether the altered migration patterns observed in MuSK MG-1 and AChR MG-1 reflect elevated IgG glycosylation as opposed to other possible modifications such as phosphorylation or ubiquitination, IgG was subjected to digestion with PNGase F or Endo S. Enzymatic digestion of MuSK MG-1 IgG with PNGase F, which non-specifically cleaves N-linked glycosylation units, resulted in a loss of the atypical IgG migration pattern. By comparison, Endo S, which cleaves N-linked glycosylation at N297 of IgG constant region, caused a shift in gel mobility in both samples, but no change in the atypical pattern (**Figure 2D**). Removal of phosphates with CIP, a phosphatase enzyme, also had no effect on migration (**Figure S3**). These results suggest that a subset of patients with MG possess atypical immunoglobulin glycosylation specifically in the Fab region, likely due to IgG-V^N-Glyc^, which appears to be a stable feature over long periods of time (3-4 years).

### MuSK and AChR human mAbs contain occupied IgG-V^N-Glyc^ sites

We had previously generated three human recombinant MuSK-specific mAbs that demonstrated *in vitro* pathogenic capacity (33, 35, 36). We found glycosylation motifs (N-X-S/T) in the variable region of all three MuSK mAbs, in either the heavy (MuSK1A and 3-28) or light chain (MuSK1B) (**Figure 3A-C**). Specifically, the motif was present in the heavy chain FWR3 of MuSK1A due to the use of IGHV1-8 where it is encoded in the germline. MuSK1B acquired the motif in the light chain (FWR1) through SHM, and the heavy chain lost the motif in the CDR2, which was present in the germline VH (IGHV4-34). MuSK3-28 acquired the motif in the heavy chain (CDR2) though SHM. We sought to test if these sites were occupied. Digestion with PNGase F reduced the MW of the heavy (MuSK1A and MuSK3-28) and light chain (MuSK2A) of the mAbs suggesting the presence of N-linked glycosylation on the antibodies (**Figure S4A-C**). We then removed these putative glycosylation sites by mutagenesis and screened all constructs for variations in migratory pattern due to MW changes. Removal of glycosylation sites led to a change in gel mobility as expected in all three MuSK mAbs, which was also consistent with site-specific occupancy (**Figure S4D-F**). Next, we performed intact mass spectrometry analysis to more precisely detect these glycosylation sites (**Figure 3A-C**). Differences in mass and mass spectra can be used to confirm the presence of IgG-V^N-Glyc^. All three MuSK autoantibodies were found to be glycosylated and the mutated variants were significantly less heterogeneous and lighter in mass by approximately 2 kDa (**Figure 3A-C**). Because N-glycans are extremely heterogenous molecular moieties, proteins containing IgG-V^N-Glyc^ have elevated mass spectra heterogeneity; these findings confirm the presence of glycosylation, and that mutations were successful in disrupting the introduction of glycosylation in all three MuSK mAbs.

**Figure 3.**
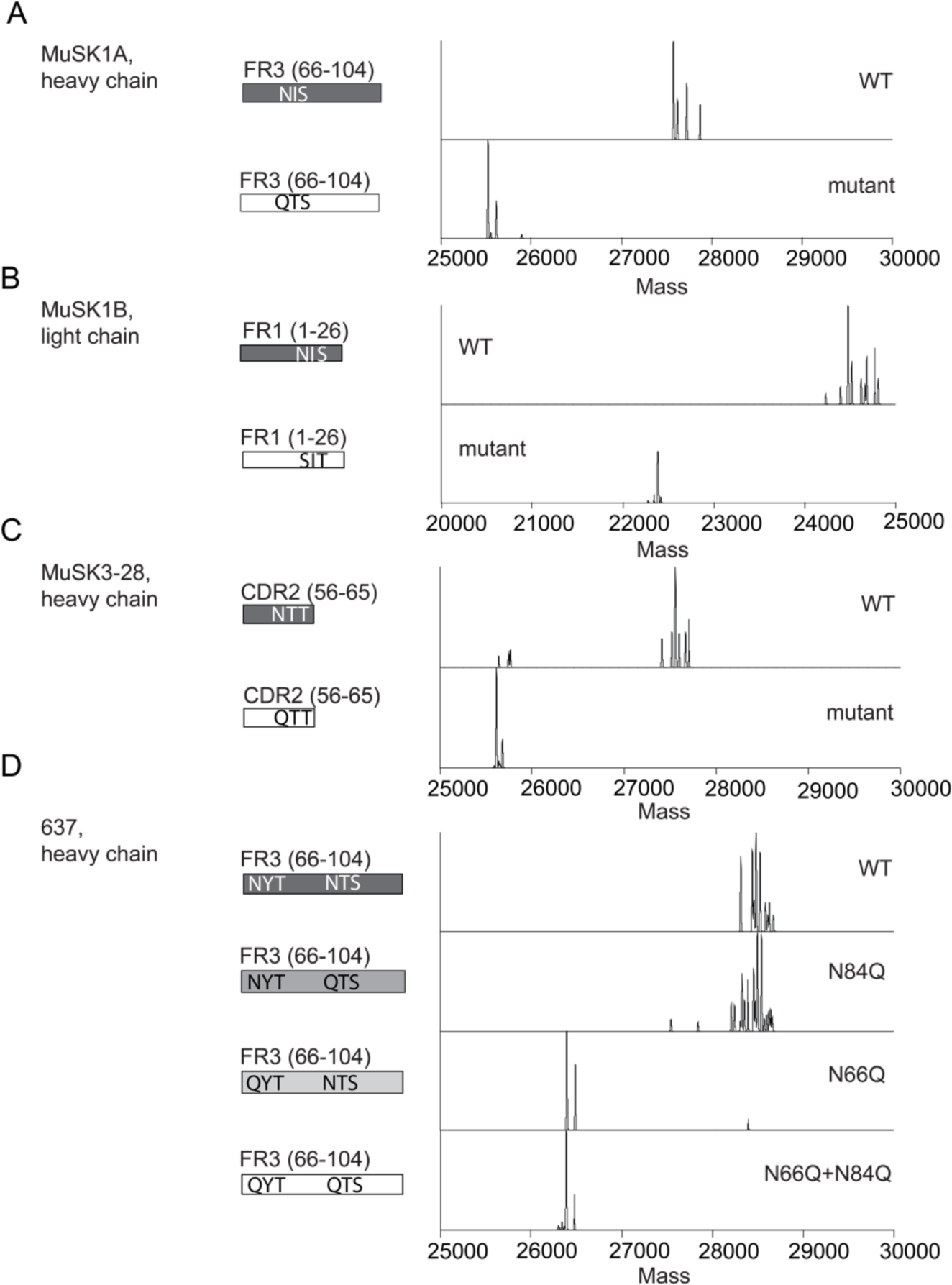
Mass spectrometry analysis of N-glycan occupancy in MuSK-specific human monoclonal antibodies. Validation of N-glycan occupancy in three patient-derived monoclonal anti-MuSK antibodies (MUSK1A, MUSK1B, MUSK3-28) and one patient-derived monoclonal anti-AChR antibody (monoclonal antibody 637). Schematic of variable regions for anti-MuSK antibodies indicating regions (CDR or FWR) and localization of putative N-linked glycosylation amino acid motifs alongside deconvoluted mass spectra of the associated constructs (labels). This is shown for MUSK1A (A), MUSK1B (B), MUSK3-28 (C) and mAb 637 (D).

Next, to extend these findings to AChR MG, we evaluated the patient-derived AChR-specific mAb 637 (37) as its sequence shows two predicted N-linked glycosylation sites (N66 and N84) within the variable region of the FR3 of heavy chain, which were acquired through SHM (**Figure 3D**). The heavy chain of mAb 637 migrated at a lower MW when treated with PNGase F in comparison to untreated mAb (**Figure S5A**). We subsequently performed mutagenesis and produced several different constructs disrupting the two predicted glycosylation sites—at N66 and N84. Mutation of N66 alone or N66 and N84 together resulted in a construct that migrated at lower MW, while mutation of only the N84 did not affect migration (**Figure S5B, C**). We further explored this result using intact mass spectrometry analysis of the Fd from the WT and three mutants (**Figure 3D**). The WT construct and the construct containing a mutation at position N84 had a complex mixture of proteoforms clustered between 28.3 – 28.8 kDa, whereas constructs containing mutations at N66 (including a mutant at both N66 and N84) were less heterogeneous, with proteoforms clustering closer to 26.4 kDa (**Figure 3D**). The lighter mass and simplified proteoform signature of variants containing a mutation at N66 suggests that N66 is the main site of glycosylation in mAb 637 and not N84. Additionally, removal of the glycosylation site N66 did not shift glycosylation to the second predicted site at N84. In summary, autoantigen-specific mAbs in MG can contain occupied N-glycosylation of their IgG variable regions.

### N-linked glycosylation in the variable region does not impact MG autoantibody binding

We then sought to test the contribution of IgG-V^N-Glyc^ to MuSK mAb binding. Given that these mAbs were previously validated for their capacity to bind AChR or MuSK in live cell-based assays (33, 35, 36), we tested the contribution of IgG-V^N-Glyc^ sites to binding using the same approach (**Figure 4A-D**). When tested over a wide range of concentrations (10 – 0.02 µg/mL), we found that loss of IgG-V^N-Glyc^ did not affect binding of anti-MuSK or anti-AChR mAbs to cognate targets.

**Figure 4.**
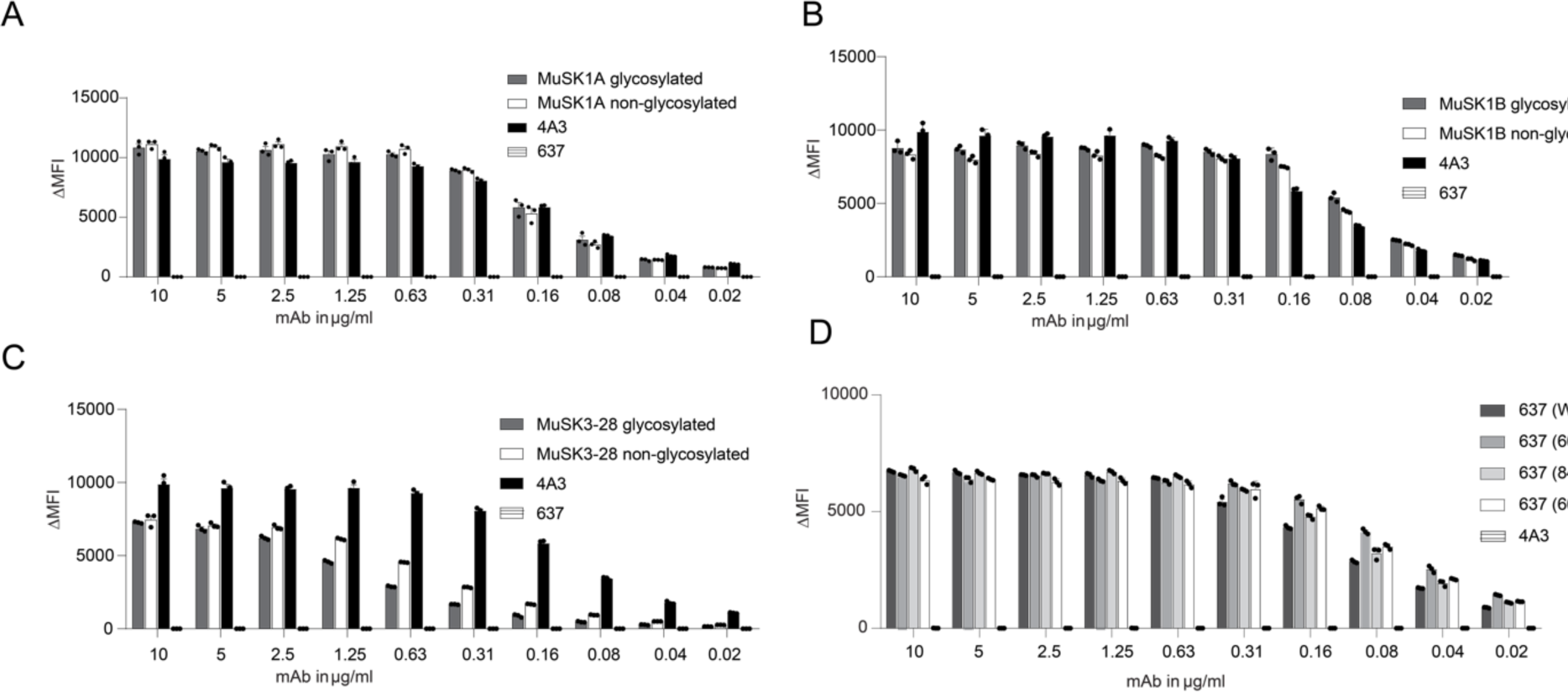
The effect of glycosylation on MuSK and AChR-specific monoclonal antibody binding properties. Antigen binding of MuSK and AChR-specific monoclonal antibodies is not affected by the presence of glycosylation sites. Wildtype MuSK and AChR mAbs and their glycovariants were tested for surface binding to MuSK or AChR on MuSK-GFP-transfected or AChR-subunit-Rapsyn-GFP-transfected HEK293T cells. (A-C) All mAb variants were analyzed in a ten two-fold dilutions series for binding to MuSK by cell-based assay (CBA). Humanized MuSK mAb 4A3 was used as the positive control and AChR-specific mAb-04 as the negative control. MuSK1A (A), MuSK1B (B) and MuSK3-28 (C) were tested. (D) All 637 mAb variants were analyzed in a ten two-fold dilutions series for binding to AChR by CBA. Humanized MuSK mAb 4A3 was used as the negative control. Each data point represents the mean value from three independent experiments, and error bars represent SDs. The ΔMFI was calculated by subtracting the signal from non-transfected cells from the signal of transfected cells.

## Discussion

Two mechanisms are thought to contribute to increased IgG-V^N-Glyc^ frequency. The first is enriched usage, at the naïve B cell stage, of the five germline V gene segments that contain N-linked glycosylation motifs (IGHV1-8, IGHV4-34, IGHV5-10, IGVL3-12 and IGVL5-37). The second mechanism is selection, during affinity maturation, of B cell clones that acquire N-linked glycosylation motifs through the SHM process. The germline encoded motifs in the three heavy chains are found in CDR2 of IGHV4-34 and in the FWR3 of IGHV1-8 and IGHV5-10. Our sequence analysis showed that these motifs, when they are acquired, are distributed throughout the variable region with the exception of FWR2. The distribution mirrors SHM patterns in that mutations accumulate preferentially in CDRs and FWR3. While replacement mutations can be observed in FWR2, our data suggest that a glycosylation motif is not tolerated in this region, suggesting that such alterations are constrained by the role of the FWRs in conserving the overall structure of the antibody. Similarly, motifs found in FWR1 and FWR4 were restricted to regions near the flexible CDR loops that they flank. These collective findings indicate that the acquisition of the motif may be driven by positive selection. It is also possible that the motifs could be selectively neutral but arise as a consequence of SHM. If so, MG repertoires could have more motifs than the HD repertoires simply by having more SHM, and the motifs could be concentrated around the CDRs due to the presence of known hotspot motifs in those regions.

SHM appears to be a major contributor to the increased frequency of the IgG-V^N-Glyc^ sites in the AChR MG patients we studied. Positive selection leading to enriched N-linked glycosylation DNA motifs has been observed in the parotid gland of patients with pSS, a structure known to contain ectopic lymphoid follicles in these patients (20). Similarly, the thymus in a subset of MG patients includes germinal centers, which are thought to contribute to the generation and maturation of AChR autoantibody producing B cells (38–40). Thus, positive selection of N-linked glycosylation DNA motifs may occur in this compartment, and support for this possibility is provided by a previous study where we found that the IgG-switched BCR sequences in MG thymus were enriched N-linked glycosylation DNA motifs (41).

Both V gene usage and the SHM process contributed to the elevated frequencies we observed in the MuSK MG patients. Defects in B cell tolerance checkpoints can skew the developing repertoire (42). Such defects are known to exist in both AChR and MuSK MG (43), and thus are likely to contribute to enrichment of the V genes containing N-linked glycosylation motifs we observed in some patients with MuSK MG. However, the accumulation of additional motifs through SHM suggests that, in the MuSK disease subtype, antigen-driven positive selection also plays a role in the conspicuously elevated IgG-V^N-Glyc^ frequency. It remains possible that this selection is an antigen-independent process.

Examples of this mechanism include interactions between glycosylated B cell receptors and lectins (21), which are thought to drive proliferation in B cell malignancies and some autoimmune diseases (20). The antigen binding by the autoantibodies we studied was not disturbed by N-linked glycosylation. Additionally, human MuSK-binding mAbs that do not include glycosylation motifs have been isolated from MG patients (34). These results indicate that selection of the IgG-V^N-Glyc^ in human MuSK autoantibodies may not have been driven by MG specific self-antigen positive selection. This is somewhat unexpected given that the IgG-V^N-Glyc^ sites could be found in regions responsible for antigen contact (CDRs). Similarly, the variable regions of anti-citrullinated protein autoantibodies (ACPA) from patients with RA are consistently glycosylated, but their binding is not influenced by the modification (18). However, other investigations suggest that binding can be modulated as a consequence of their presence (5, 16, 17). These collective findings suggest that an autoantigen-independent selection mechanism may influence the IgG-V^N-Glyc^ motif frequency in the autoimmune repertoire in some, but not all, autoimmune diseases.

Our proteomic analysis of the serum-derived IgG from only two of the study subjects (one from each of the AChR and MuSK MG cohorts) showed a higher molecular weight band in the electrophoresis studies. These findings indicate that the serum IgG repertoire may not be well reflected by the circulating IgG B cell repertoire that we sequenced, which has been previously suggested (44). Rather, these findings may reflect that much of the circulating IgG is derived from long-lived plasma cells residing in the bone marrow. Furthermore, other investigations (14, 16) that showed the presence of IgG-V^N-Glyc^ by electrophoresis in human autoimmunity, focused on specifically enriched autoantibodies rather than total circulating IgG, which was the focus of our study.

One of two possible glycosylation sites in an autoantibody known to be specific for AChR, mAb 637, was shown to be unoccupied. We speculate that this is indicative of context specific N-glycosylation (local amino acid sequence containing the motif or cellular environment) or inherent selectivity for one site over the other possibly due to conformation or solvent accessibility. Nevertheless, this site did not appear to contribute to binding activity, similar to our observations obtained by testing the MuSK mAbs. We recognize, as a study limitation, that the *in vitro* expression of these mAbs may not emulate the glycosylation occupancy *in vivo*. We did use a mammalian expression system (human embryonic kidney cells), to achieve the best approximation of the *in vivo* status, and we experimentally confirmed occupancy for the antigen binding studies. It remains to be investigated whether variables such as the stage of B cell activation or tissue residence could alter the occupancy.

Finally, a consensus on the function of IgG-V^N-Glyc^ in health or disease is unclear. Several possibilities have been described, including perturbation of antibody-antigen interactions (binding affinity, specificity), altered metabolism of B cells or IgG *in vivo* (half-life, clearance), mis-localization of IgG to host tissue, redemption of autoreactive B cells, and inappropriate selection/expansion of autoreactive B cells in germinal centers (1, 2, 45). Elimination of these motifs or removal of the glycosyl moiety itself have been observed to impair antigen binding—such as in anti-adalimumab/infliximab antibodies derived from patients treated for RA (5). However other studies have suggested a more nuanced picture; a study of anti-CCP (cyclic citrullinated protein) autoantibodies showed no contribution of N-linked glycans to binding (18). Here we show unequivocal evidence that the presence of N-linked glycans is not required for binding in the case of four MG autoantibodies. This appears to agree with the majority of studies published regarding the role of IgG-V^N-Glyc^ on antigen binding.

In summary, IgG-V^N-Glyc^ is elevated in a subset of patients AChR and MuSK MG. These findings are consistent with those of a previous study (46) that showed elevated (albeit not statistically significant) V region N-glycosylation sites in the B cell repertoire of MG patients compared to healthy controls. Our findings extend this molecular phenotype beyond RA, pSS, SLE, and AAV. IgG-V^N-Glyc^ does not affect AChR or MuSK autoantibody binding. We speculate an elevation in N-linked glycosylation motifs containing V gene sequences may be driven by the presence of dysregulated germinal centers that contribute to B cell selection defects observed in the disease. Our findings contribute to efforts to understand the basic biology of IgG-V^N-Glyc^ and its association with disease.

## Methods

### Patient selection

This study was approved by Yale University’s Institutional Review Board (clinicaltrials.gov || NCT03792659). Informed consent was received from all participating patients prior to inclusion in this study. Peripheral blood was collected from MG subjects at the Yale Myasthenia Gravis Clinic, New Haven, Connecticut, USA (47). Informed consent was received from all participating patients prior to inclusion in this study. A MuSK MG cohort (N=3) was defined using BCR repertoire sequencing derived from our previous study (48). Another n = 10 AChR MG and n = 9 heathy control subjects were selected for BCR based adaptive immune receptor repertoire-sequencing or AIRR-Seq using PBMC derived RNA for this study. In total, serum samples from all 3 MuSK MG subjects and 9 AChR MG subjects (8 overlapping with paired AIRR-Seq) cohort were also investigated for the presence of VH gene glycosylation-specific signatures in the serum. With the exception of patient 4, AChR MG patients had not received any immunotherapy or prednisone prior to sample collection. For patients with MuSK MG diagnoses and patient AChR MG-1 with an AChR MG diagnosis, longitudinal serum samples were collected. A patient with Rheumatoid Arthritis (RA) (n = 1) was enrolled in a research study at University of California San Francisco (UCSF) for pathogen and autoantibody detection.

### Protein electrophoresis and immunoblotting

For the initial screening processes patient sera samples were diluted 1:1 with 2X Storage Buffer (2X PBS, 20mM HEPES, 0.04% Sodium Azide, 20% Glycerol). The IgG from human sera or MuSK-specific mAb 4A3 and AChR-specific mAb 637 were captured with AG beads (Thermo Fisher) and then eluted by boiling at 95 degrees in 2X Laemmli buffer (with 10% beta-mercaptoethanol). For immunoblotting the gel was transferred to 0.45-micron nitrocellulose membrane and blotted with secondary anti-human IgG conjugated to IR800 dye (LICOR, Cat). Nitrocellulose blots were imaged with LICOR scanner and analyzed qualitatively by eye for presence of altered migration patterns in IgG. Potential IgG heavy chain migration phenotypes were qualitatively called by an experimenter blind to experimental conditions. For the three MuSK-specific (mAb MuSK1A, MuSK1B and MuSK3-28) and AChR-specific mAb 637 Mini-PROTEAN® TGX Stain-Free™ Precast Gels (Biorad) and Laemmli Sample Buffer (Biorad) were used for SDS-page. Prior to electrophoresis the proteins were reduced with 0.1 M DTT (Thermo Fisher Scientific) and heat denatured at 95°C for 5 min. After electrophoresis the gel was stained with Coomassie blue solution. Bands were visualized with the ChemiDoc™ Touch Imaging System (Biorad). Enzymatic assays for PNGase F and Endo S were performed according to manufacturer’s instructions (NEB). The effect of the enzymatic assays was either analyzed by Coomassie staining or immunoblotting.

### BCR library preparation, pre-processing and analysis

First, RNA was isolated from frozen peripheral blood mononuclear cells using the RNAEasy Mini kit (Qiagen) per manufacturer’s instructions. Bulk libraries were prepared from RNA using reagents from New England Biolabs as part of the NEBNext® Immune Sequencing Kit as described previously (41, 42). Briefly, cDNA was reverse-transcribed by a template-switch reaction to add a 17-nucleotide unique molecular identifier (UMI) to the 5’ end with streptavidin magnetic bead purification. This was then followed by two rounds of PCR; the first round enriched for immunoglobulin sequences using IGHA, IGHD, IGHE, IGHG, and IGHM-specific 3’ primers and added a 5’ index primer. Libraries were purified with AMPure XP beads (Beckman) after which another round of PCR added Illumina P5 Adaptor sequences to each amplicon. The number of cycles selected based on quantitative PCR to avoid the plateau phase. Libraries were then purified again with AMPure beads. Libraries were pooled in equimolar libraries and sequenced by 325 cycles for read 1 and 275 cycles for read 2 using paired-end sequencing with a 20% PhiX spike on the Illumina MiSeq platform according to manufacturer’s recommendations.

Processing and analysis of bulk B cell receptor sequences was carried out using tools from the Immcantation framework as done previously (49). Preprocessing was performed using pRESTO. Briefly, sequences with a phred score below 20 were removed and only those that contained constant region and template switch sequences were preserved. UMI sequences were then grouped and consensus sequences were constructed for each group and assembled into V(D)J sequences in a two-step process involving an analysis of overlapping sequences (<8 nucleotides) or alignment against the IMGT (the international ImMunoGeneTics information system®) IGHV reference (IMGT/GENE-DB v3.1.19; retrieved December 1, 2019) if no significant overlap was found. Isotypes were assigned by local alignment of the 3’ end of the V(D)J sequence to constant region sequences. Duplicate sequences were removed and only V(D)J sequences reconstructed from more than 1 amplicon were preserved. Primer sequences used for this analysis are available at: https://bitbucket.org/kleinstein/immcantation.

V(D)J germline genes were assigned to reconstructed V(D)J sequences using IgBLAST v.1.14.0 also using the December 1, 2019 version of the IMGT gene database for both bulk and single cell repertoires (50). V(D)J sequences with IGH associated V and J genes were then selected for further analysis and non-functional sequences were removed. Germline sequences were reconstructed for each V(D)J sequence with D segment and N/P regions masked (with Ns) using the CreateGermlines.py function within Change-O v1.0.0(51). VH gene nucleotides up to IMGT position 312 were translated from both the aligned sequence and germline reconstructed V(D)J sequence using BioPython v1.75. To quantify the frequency of N-X-S/T glycosylation motifs, matches to the regular expression pattern “N[^P][S,T]” were quantified for each translated sequence, including for translated CDR and FWR fragments of the VH gene sequence (defined by IMGT coordinates) (4). N-X-S/T glycosylation motifs in the CDR3 and FWR4 regions were similarly quantified separately and included for CDR and FWR distribution analyses.

To build the lineage tree in **Supplemental Figure 1**, B cells were first clustered into clones by partitioning based on common IGHV gene annotations, IGHJ gene annotations, and junction lengths. Within these groups, sequences differing from one another by a length normalized Hamming distance of 0.2 within the junction region were defined as clones by single-linkage clustering using Change-O v.1.0.1(51). The Hamming distance threshold was determined by manual inspection of the distance to the nearest sequence neighbor plot using SHazaM v1.0.2(52). Phylogenetic tree topology and branch lengths of an illustrative clonal lineage were estimated using the HLP19 model in IgPhyML v1.1.3 and visualized using ggtree v2.0.4 and custom R scripts(53, 54).

### Mass Spectrometry

High-resolution mass spectrometry (HRMS) was employed to confirm the change in glycosylation status between wild type and mutated variants. In order to reduce complexity at the intact mass level a “middle-down” approach was utilized (55, 56). Intact antibodies were incubated with IdeS protease, followed by reduction of disulfide bonds. This workflow is well known to break down antibodies into three ∼25 kDa subunits – LC, Fc, and Fd – and thereby separate disease-associated glycosylation within the variable region from standard glycosylation in the constant region. The heavy chain variable region (VH) is located within the Fd subunit. Purified mAb (2 mg/mL in PBS) was treated with 1 unit of IdeS protease (Promega) per 1 µg of mAb, and the sample was incubated at 37°C in a shaking incubator for 1.5 hours. The digested sample was then diluted into 6 M guanidinium chloride to a final IgG concentration of 1 mg/mL, and Tris(2-carboxyethyl) phosphine hydrochloride (TCEP-HCl) was added for a final concentration of 30 mM. The sample was incubated at 37°C in a shaking incubator for 1.5 hours, then the reaction was quenched by the addition of trifluoroacetic acid (final TFA concentration of 0.1% v/v). The sample was desalted by buffer exchange into LC-MS buffer (5 rounds of buffer exchange with an Amicon Ultra-0.5 mL centrifugal filter unit, 10 kDa MWCO).

The digested and reduced antibody species were further desalted and separated on a monolithic C4 column (RP-5H, 100 mm, 0.5 mm i.d., Thermo Scientific) with an Ultimate 3000 RSLCnano system (Thermo Scientific) using a binary gradient. The gradient utilized solvent A: 95% water, 5% acetonitrile, and 0.2% formic acid, and solvent B: 5% water, 95% acetonitrile, and 0.2% formic acid.

Data were acquired on a Q Exactive HF instrument with an attached HESI source (sheath gas = 10, auxiliary gas = 2, spare gas = 2, spray voltage = 3500 V, S-lens RF level = 65). MS1 acquisition used a scan range window of 400 to 2,000 m/z with 1 microscan and an AGC target of 1e6, at a resolution of 15,000.

### Site directed mutagenesis of glycosylation site

Glycosylation sites (N-X-S/T) present in the V regions of the monoclonal antibodies were removed by mutating the asparagine (N) either to a glutamine (Q) or a serine (S). This was performed with Q5® Site-Directed Mutagenesis Kit (NEB) according to manufacturer’s instructions. The primers were designed with NEBaseChanger. Sequences of all expression plasmids were verified by Sanger sequencing.

### Recombinant expression of human monoclonal antibodies (mAbs)

The mAbs were produced as previously described (Takata et al., 2019). Briefly, HEK293A cells were transfected with equal amounts of the heavy and the corresponding light chain plasmid using linear PEI (Polysciences Cat# 23966). The media was changed after 24 h to BASAL media (50% DMEM 12430, 50% RPMI 1640, 1% antibiotic/antimycotic, 1% Na-pyruvate, 1% Nutridoma). After 6 days the supernatant was harvested and Protein G Sepharose® 4 Fast Flow beads (GE Healthcare) were used for antibody purification.

### Live cell-based autoantibody assay

Cell-based assays for detection of AChR or MuSK antibody binding were performed as we have previously described(57). Briefly, the cDNA encoding human AChR α, β, δ, ε-subunits and rapsyn-GFP were each cloned into pcDNA3.1-hygro plasmid vectors (Invitrogen, CA) and cDNA encoding human full-length MuSK was cloned into pIRES2-EGFP plasmid vector (Clontech). AChR and MuSK vectors were kindly provided by Drs. D. Beeson and A. Vincent of the University of Oxford. HEK293T (ATCC^®^ CRL3216™) cells were transfected with either MuSK-GFP, or the AChR domains together with rapsyn-GFP. On the day of the CBA, the mAbs were added to the transfected cells in a dilution series (10 – 0.02 μg/ml). The binding of each mAb was detected with Alexa Fluor®-conjugated AffiniPure Rabbit Anti-Human IgG, Fcγ (309-605-008, Jackson Immunoresearch) on a BD LSRFortessa® (BD Biosciences). FlowJo software (FlowJo, LLC) was used for analysis.

### Statistics

R v4.0.3 was used for all statistical analysis. Data frame handling and plotting was performed using functions from the tidyverse v1.3.0 in R and pandas v0.24.2 in python v3.7.5. A significance threshold of <0.05 was used and shown on plots with a single asterisk; double asterisks correspond to a p <0.01 and triple asterisks correspond to a p<0.001. Unpaired one-tailed Wilcoxon tests were used for comparisons with healthy controls in repertoire analysis; the alternative hypothesis was that the average count of glycosylation motifs for each V(D)J sequence in MG BCR repertoires would be higher.

## Acknowledgments

The authors thank Karen Boss for expert copy editing and proofreading, Dr. Bailey Munro-Sheldon for verifying autoantibody titers, and Charlotte A. Gurley for assisting with manuscript and reference formatting.

---

**Supplemental Figure 1.**
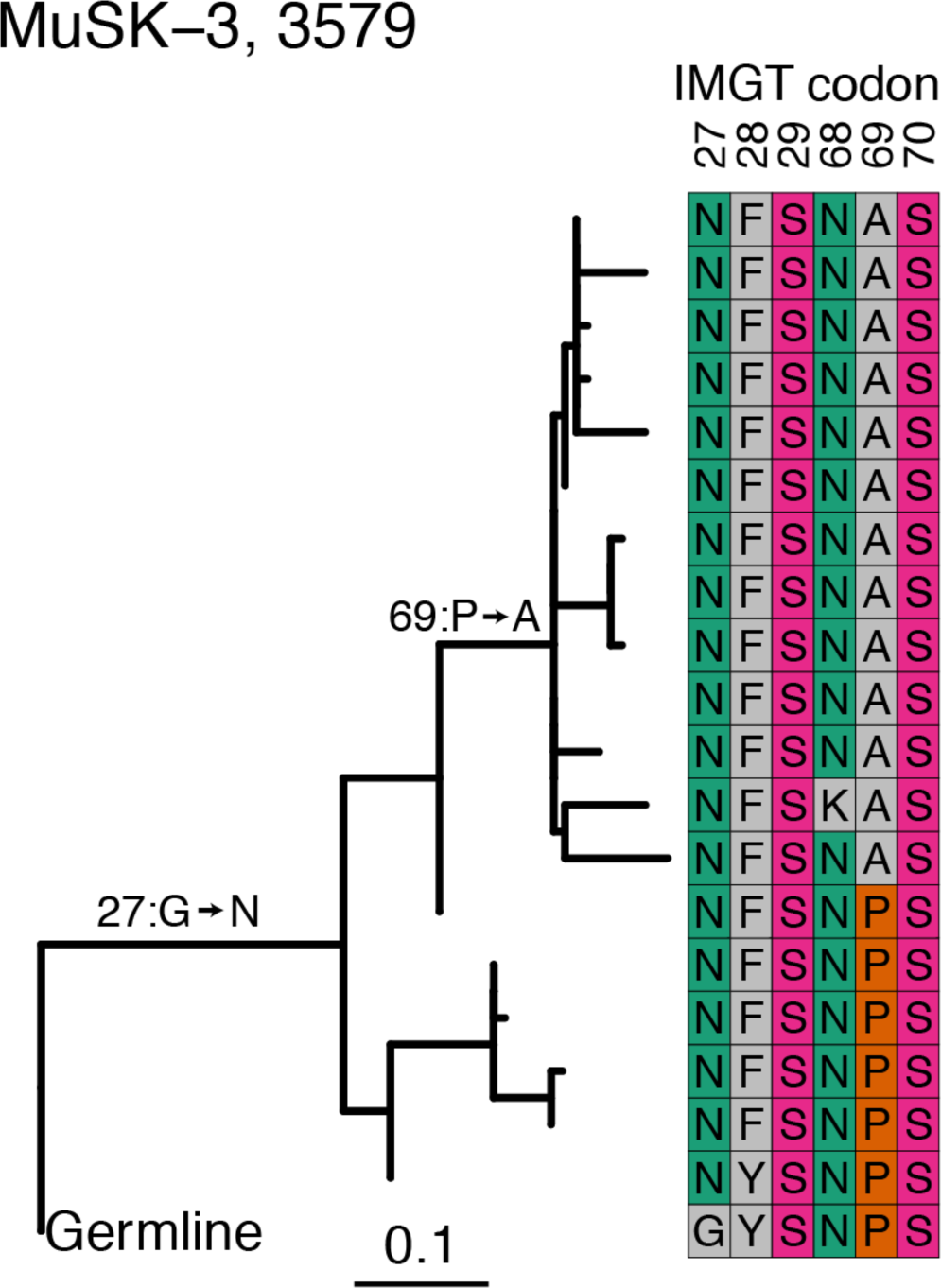
Clonal lineage V^N-Glyc^ motif acquisition through somatic hypermutation. An illustrative example showing a maximum likelihood tree corresponding to a clone that acquired two unique N-X-S/T motifs during the somatic hypermutation process. Edge lengths are quantified based on number of intervening somatic hypermutations per codon between nodes per the scale. This example shows clone 3579 from patient MuSK MG-3, which uses IGHV4-38-2. The acquisition of the first N-X-S/T motif in the CDR1 (codon 27 using IMGT numbering) occurs early in the clonal development and is maintained throughout the lineage. A second motif is acquired in the FWR3 (codon 68 using IMGT numbering), close to the CDR2.

**Supplemental Figure 2.**
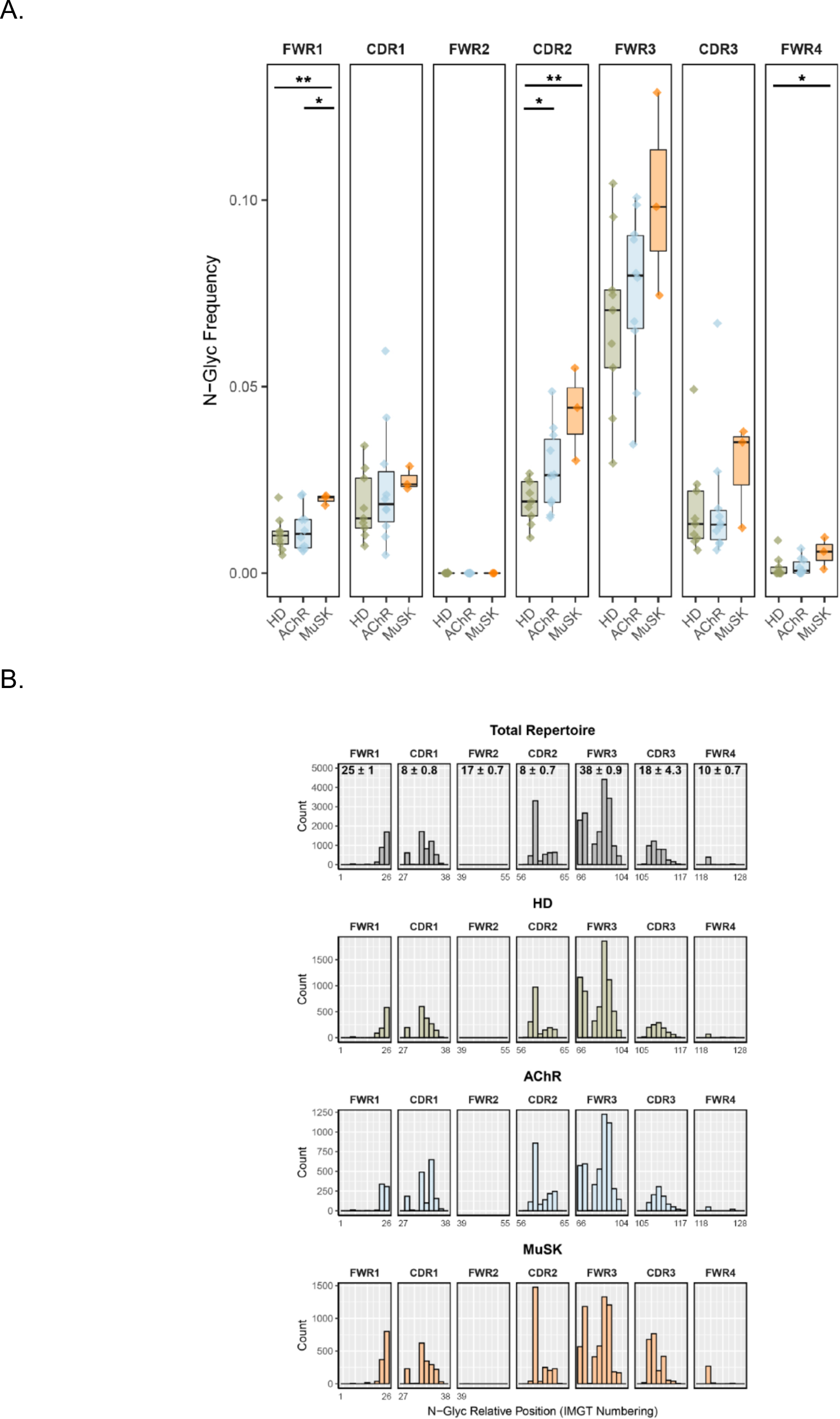
Distribution of IgG isotype-specific V^N-Glyc^ sites in the BCR. B cell receptor sequence analysis showing elevated frequency of N-linked glycosylation motifs in HD, AChR MG and MuSK MG repertoires across CDRs and FWRs (**A**). Frequency of N-linked glycosylation motifs (N-X-S/T, average number of sites per V(D)J sequence) in each V gene region of IgG sequences is shown. Title over each panel specifies the region that was searched. A significance threshold of p <0.05 was used and shown on plots with a single asterisk; double asterisks correspond to a p <0.01. Histogram (**B**) showing positional distribution of N-linked glycosylation motifs within the FWRs and CDRs. The average length (AA) and standard deviation of each region is indicated in the panels on the top row.

**Supplemental Figure 3.**
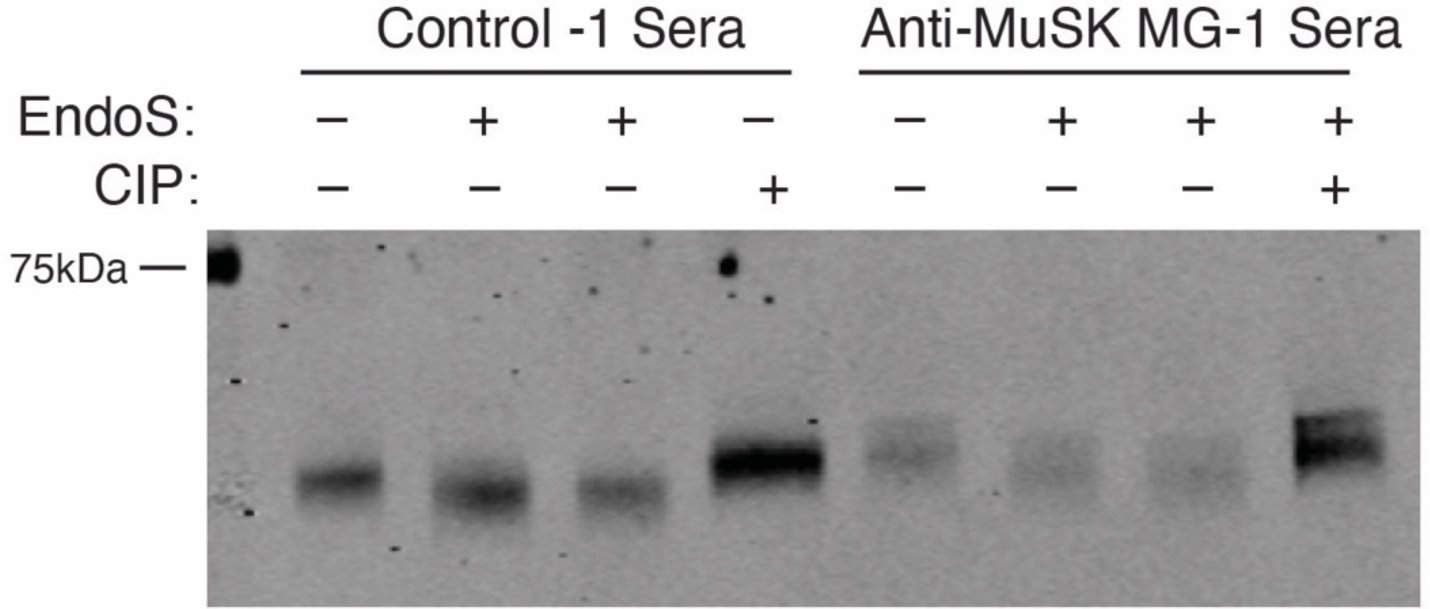
Additional enzymatic digestions of MG serum-derived IgG to deduce the molecular basis of altered heavy chain migration pattern. Treatment of IgG with Endoglycosidase (EndoS) or Calf intestinal phosphates CIP had no effect on migration as indicated by presence of double band in all conditions in MG. EndoS cleaves the chitobiose core of N-linked glycans, leaving the primary N-acetylglucosamine linked to Asparagine. CIP catalyzes dephosphorylation.

**Supplemental Figure 4.**
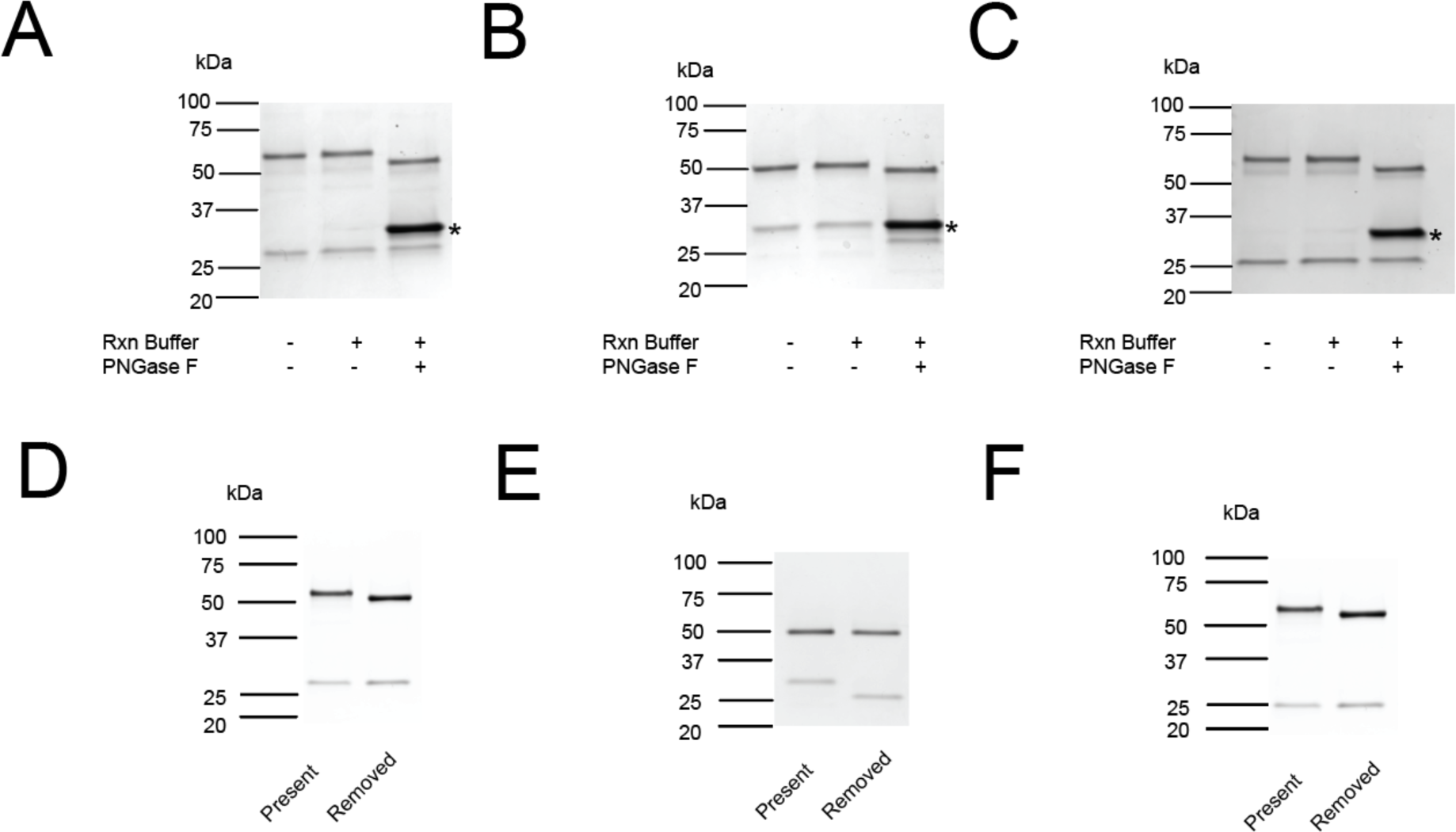
IgG migration patterns of heavy and light chains from wildtype mAbs following removal of N-glycans. MuSK1A (A), MuSK1B (B) and MuSK3-28 (C) were treated with the enzyme PNGase F as indicated. Subsequently, the proteins were separated by SDS-PAGE and detected by Coomassie staining. Asterisk marks PNGase F enzyme (A-C). (D-F) All three mature MuSK mAb contained glycosylation motifs. The motifs were removed through mutagenesis and the proteins tested by Coomassie staining for consecutive change of molecular weight (MW). Present corresponds to the WT construct while Removed refers to the construct with the N-glycan site mutated (D-F).

**Supplemental Figure 5.**
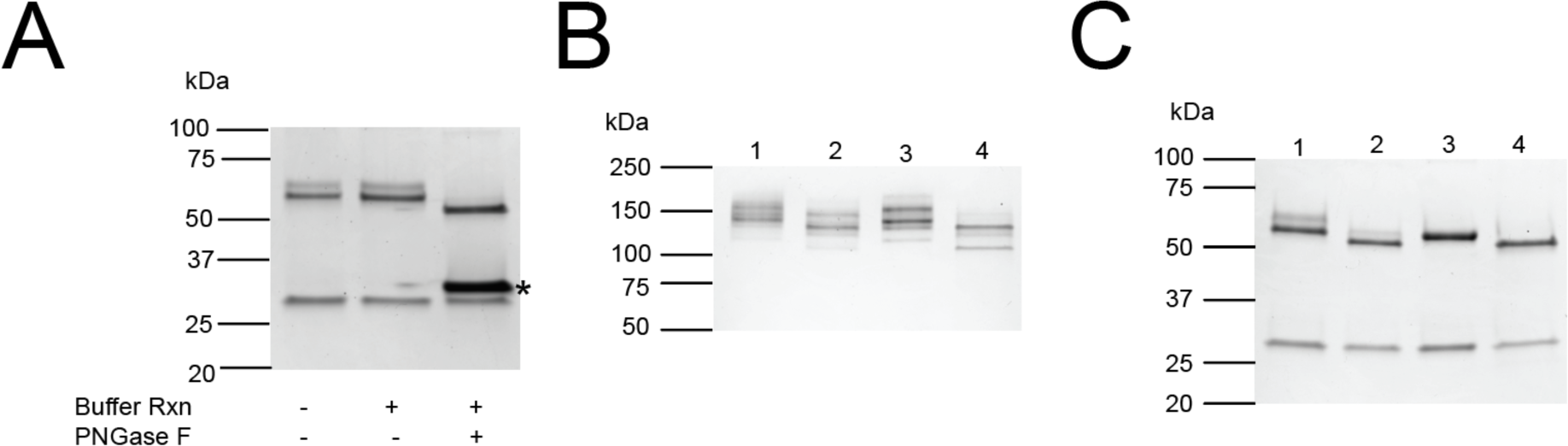
Proteomic analysis of glycosylation for AChR-specific mAb 637. (A) The mAb 637 was treated with the enzyme PNGase F as indicated. Subsequently, the proteins were separated by SDS-PAGE and detected by Coomassie staining. (B) (C) The generated knockout constructs were tested by Coomassie staining for consecutive change of molecular weight (MW). The constructs were either loaded untreated (B) or reduced by DTT and boiled at 95°C for 5 min (C). Asterisk marks PNGase F enzyme in (A). In (B) and (C), the lanes correspond to the following: Lane 1: 637 (WT); Lane 2: 637 (66Q); Lane 3: 637 (84Q); Lane 4: 637 (66Q+84Q).

**Supplemental Table 1.**
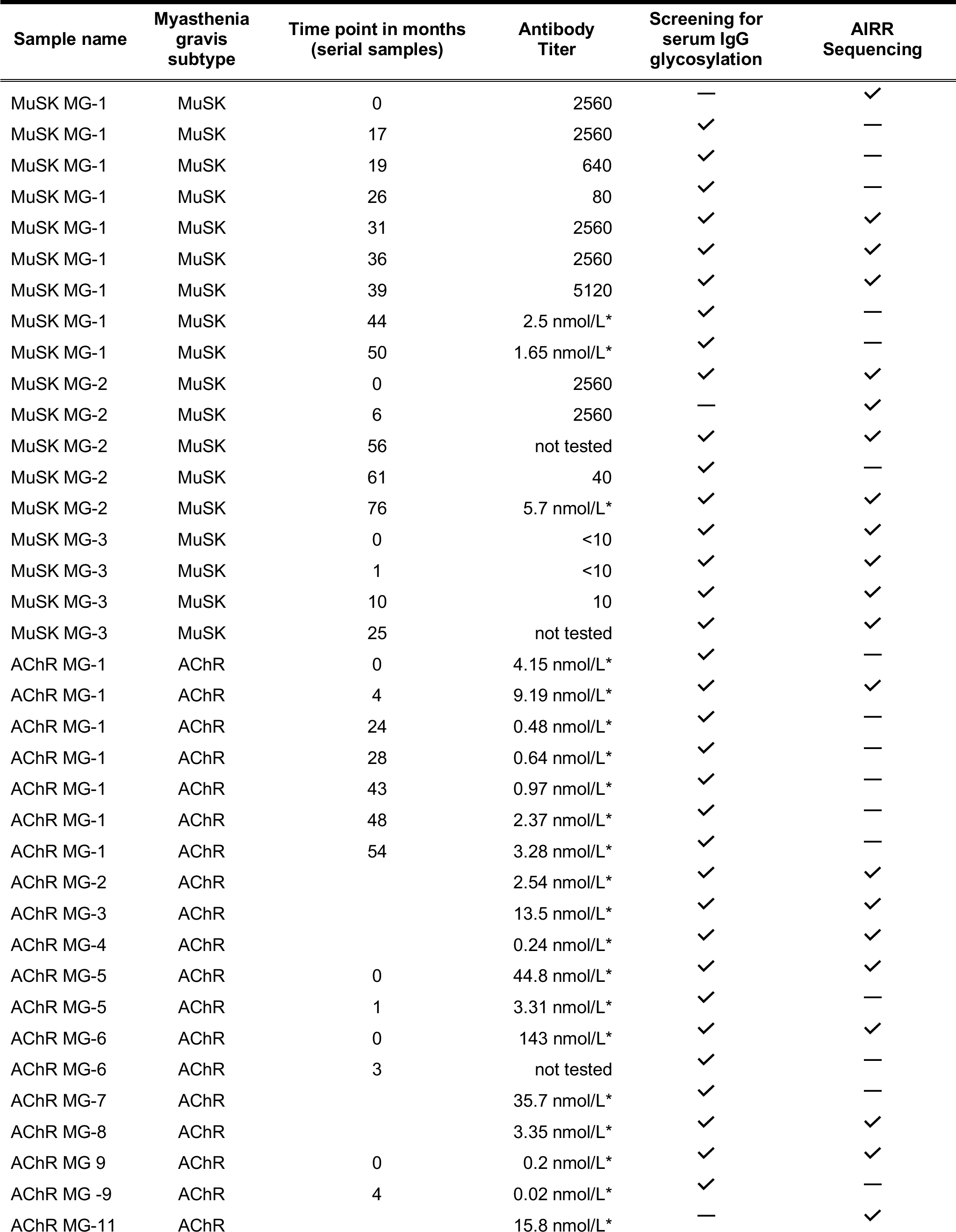
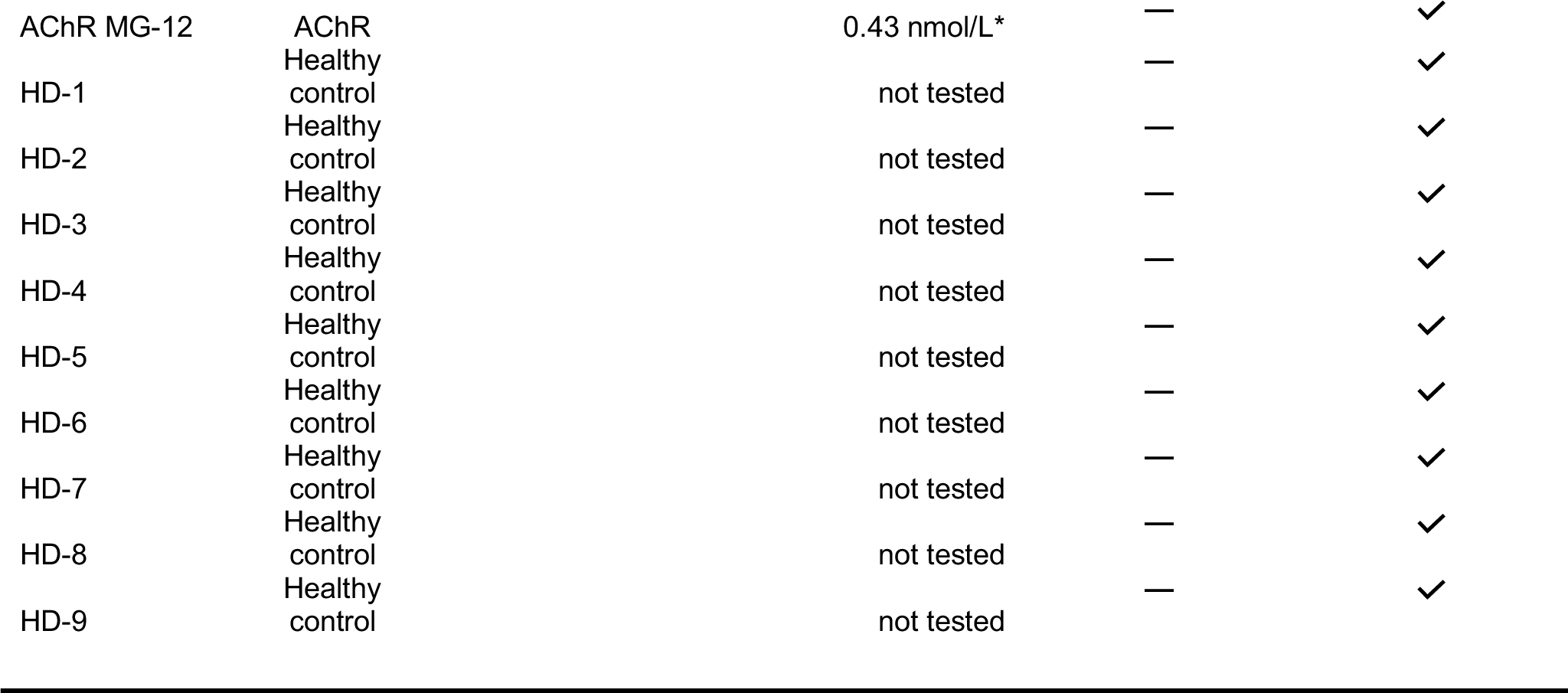
Characteristics and analysis status of study subjects. Myasthenia gravis subtype, time points of collected serial samples and subtype-specific autoantibody titer/concentration of each study specimen. The reference range for positivity varies according to the measuring facility. For samples measured by Athena Diagnostics the titer range is negative for <1:10, borderline for 1:10 and positive for >1:20. The cut off for negativity for samples measured at Mayo Clinic Laboratory is ≤ 0.02 nmol/L. Samples measured at Mayo Clinic Laboratory are indicated by an (*). Analysis status for serum IgG glycosylation or AIRR sequencing data is indicated by (✓) for available and (**—)** for not performed. The time point 0 of each serial sample is normalized to indicate the first sample in the series.

## References

1. van de Bovenkamp, F. S., L. Hafkenscheid, T. Rispens, and Y. Rombouts. 2016. The Emerging Importance of IgG Fab Glycosylation in Immunity. J Immunol 196: 1435–1441.

2. Kanyavuz, A., A. Marey-Jarossay, S. Lacroix-Desmazes, and J. D. Dimitrov. 2019. Breaking the law: unconventional strategies for antibody diversification. Nat Rev Immunol 19: 355–368.

3. Dunn-Walters, D., L. Boursier, and J. Spencer. 2000. Effect of somatic hypermutation on potential N-glycosylation sites in human immunoglobulin heavy chain variable regions. Mol Immunol 37: 107–113.

4. Lefranc, M. P. 2001. IMGT, the international ImMunoGeneTics database. Nucleic Acids Res 29: 207–209.

5. van de Bovenkamp, F. S., N. I. L. Derksen, P. Ooijevaar-de Heer, K. A. van Schie, S. Kruithof, M. A. Berkowska, C. E. van der Schoot, I. J. H, M. van der Burg, A. Gils, L. Hafkenscheid, R. E. M. Toes, Y. Rombouts, R. Plomp, M. Wuhrer, S. M. van Ham, G. Vidarsson, and T. Rispens. 2018. Adaptive antibody diversification through N-linked glycosylation of the immunoglobulin variable region. Proc Natl Acad Sci U S A 115: 1901–1906.

6. Zhu, D., H. McCarthy, C. H. Ottensmeier, P. Johnson, T. J. Hamblin, and F. K. Stevenson. 2002. Acquisition of potential N-glycosylation sites in the immunoglobulin variable region by somatic mutation is a distinctive feature of follicular lymphoma. Blood 99: 2562–2568.

7. Radcliffe, C. M., J. N. Arnold, D. M. Suter, M. R. Wormald, D. J. Harvey, L. Royle, Y. Mimura, Y. Kimura, R. B. Sim, S. Inoges, M. Rodriguez-Calvillo, N. Zabalegui, A. L. de Cerio, K. N. Potter, C. I. Mockridge, R. A. Dwek, M. Bendandi, P. M. Rudd, and F. K. Stevenson. 2007. Human follicular lymphoma cells contain oligomannose glycans in the antigen-binding site of the B-cell receptor. J Biol Chem 282: 7405–7415.

8. Odabashian, M., E. Carlotti, S. Araf, J. Okosun, F. Spada, J. G. Gribben, F. Forconi, F. K. Stevenson, M. Calaminici, and S. Krysov. 2020. IGHV sequencing reveals acquired N-glycosylation sites as a clonal and stable event during follicular lymphoma evolution. Blood 135: 834–844.

9. Koning, M. T., E. Quinten, W. H. Zoutman, S. M. Kielbasa, H. Mei, C. A. M. van Bergen, P. Jansen, R. D. Vergroesen, R. Willemze, M. H. Vermeer, C. P. Tensen, and H. Veelken. 2019. Acquired N-Linked Glycosylation Motifs in B-Cell Receptors of Primary Cutaneous B-Cell Lymphoma and the Normal B-Cell Repertoire. J Invest Dermatol 139: 2195–2203.

10. Zabalegui, N., A. L. de Cerio, S. Inoges, M. Rodriguez-Calvillo, J. Perez-Calvo, M. Hernandez, J. Garcia-Foncillas, S. Martin-Algarra, E. Rocha, and M. Bendandi. 2004. Acquired potential N-glycosylation sites within the tumor-specific immunoglobulin heavy chains of B-cell malignancies. Haematologica 89: 541–546.

11. Visser, A., N. Hamza, F. G. M. Kroese, and N. A. Bos. 2018. Acquiring new N-glycosylation sites in variable regions of immunoglobulin genes by somatic hypermutation is a common feature of autoimmune diseases. Ann Rheum Dis 77: e69.

12. Holland, M., H. Yagi, N. Takahashi, K. Kato, C. O. Savage, D. M. Goodall, and R. Jefferis. 2006. Differential glycosylation of polyclonal IgG, IgG-Fc and IgG-Fab isolated from the sera of patients with ANCA-associated systemic vasculitis. Biochim Biophys Acta 1760: 669–677.

13. Xu, P. C., S. J. Gou, X. W. Yang, Z. Cui, X. Y. Jia, M. Chen, and M. H. Zhao. 2012. Influence of variable domain glycosylation on anti-neutrophil cytoplasmic autoantibodies and anti-glomerular basement membrane autoantibodies. BMC Immunol 13: 10.

14. Lardinois, O. M., L. J. Deterding, J. J. Hess, C. J. Poulton, C. D. Henderson, J. C. Jennette, P. H. Nachman, and R. J. Falk. 2019. Immunoglobulins G from patients with ANCA-associated vasculitis are atypically glycosylated in both the Fc and Fab regions and the relation to disease activity. PLoS One 14: e0213215.

15. Youings, A., S. C. Chang, R. A. Dwek, and I. G. Scragg. 1996. Site-specific glycosylation of human immunoglobulin G is altered in four rheumatoid arthritis patients. Biochem J 314 ( Pt 2): 621–630.

16. Rombouts, Y., A. Willemze, J. J. van Beers, J. Shi, P. F. Kerkman, L. van Toorn, G. M. Janssen, A. Zaldumbide, R. C. Hoeben, G. J. Pruijn, A. M. Deelder, G. Wolbink, T. Rispens, P. A. van Veelen, T. W. Huizinga, M. Wuhrer, L. A. Trouw, H. U. Scherer, and R. E. Toes. 2016. Extensive glycosylation of ACPA-IgG variable domains modulates binding to citrullinated antigens in rheumatoid arthritis. Ann Rheum Dis 75: 578–585.

17. Hafkenscheid, L., A. Bondt, H. U. Scherer, T. W. Huizinga, M. Wuhrer, R. E. Toes, and Y. Rombouts. 2017. Structural Analysis of Variable Domain Glycosylation of Anti-Citrullinated Protein Antibodies in Rheumatoid Arthritis Reveals the Presence of Highly Sialylated Glycans. Mol Cell Proteomics 16: 278–287.

18. Lloyd, K. A., J. Steen, K. Amara, P. J. Titcombe, L. Israelsson, S. L. Lundstrom, D. Zhou, R. A. Zubarev, E. Reed, L. Piccoli, C. Gabay, A. Lanzavecchia, D. Baeten, K. Lundberg, D. L. Mueller, L. Klareskog, V. Malmstrom, and C. Gronwall. 2018. Variable domain N-linked glycosylation and negative surface charge are key features of monoclonal ACPA: Implications for B-cell selection. Eur J Immunol 48: 1030–1045.

19. Visser, A., M. E. Doorenspleet, N. de Vries, F. K. L. Spijkervet, A. Vissink, R. J. Bende, H. Bootsma, F. G. M. Kroese, and N. A. Bos. 2018. Acquisition of N-Glycosylation Sites in Immunoglobulin Heavy Chain Genes During Local Expansion in Parotid Salivary Glands of Primary Sjogren Patients. Front Immunol 9: 491.

20. Hamza, N., U. Hershberg, C. G. Kallenberg, A. Vissink, F. K. Spijkervet, H. Bootsma, F. G. Kroese, and N. A. Bos. 2015. Ig gene analysis reveals altered selective pressures on Ig-producing cells in parotid glands of primary Sjogren’s syndrome patients. J Immunol 194: 514–521.

21. Coelho, V., S. Krysov, A. M. Ghaemmaghami, M. Emara, K. N. Potter, P. Johnson, G. Packham, L. Martinez-Pomares, and F. K. Stevenson. 2010. Glycosylation of surface Ig creates a functional bridge between human follicular lymphoma and microenvironmental lectins. Proc Natl Acad Sci U S A 107: 18587–18592.

22. Gilhus, N. E., G. O. Skeie, F. Romi, K. Lazaridis, P. Zisimopoulou, and S. Tzartos. 2016. Myasthenia gravis - autoantibody characteristics and their implications for therapy. Nat Rev Neurol 12: 259–268.

23. Vincent, A. 2002. Unravelling the pathogenesis of myasthenia gravis. Nat Rev Immunol 2: 797–804.

24. Yi, J. S., J. T. Guptill, P. Stathopoulos, R. J. Nowak, and K. C. O’Connor. 2018. B cells in the pathophysiology of myasthenia gravis. Muscle & nerve 57: 172–184.

25. Hoch, W., J. McConville, S. Helms, J. Newsom-Davis, A. Melms, and A. Vincent. 2001. Auto-antibodies to the receptor tyrosine kinase MuSK in patients with myasthenia gravis without acetylcholine receptor antibodies. Nat Med 7: 365–368.

26. McConville, J., M. E. Farrugia, D. Beeson, U. Kishore, R. Metcalfe, J. Newsom-Davis, and A. Vincent. 2004. Detection and characterization of MuSK antibodies in seronegative myasthenia gravis. Ann Neurol 55: 580–584.

27. Fichtner, M. L., R. Jiang, A. Bourke, R. J. Nowak, and K. C. O’Connor. 2020. Autoimmune Pathology in Myasthenia Gravis Disease Subtypes Is Governed by Divergent Mechanisms of Immunopathology. Front Immunol 11: 776.

28. Drachman, D. B., R. N. Adams, L. F. Josifek, and S. G. Self. 1982. Functional activities of autoantibodies to acetylcholine receptors and the clinical severity of myasthenia gravis. N Engl J Med 307: 769–775.

29. Drachman, D. B., C. W. Angus, R. N. Adams, J. D. Michelson, and G. J. Hoffman. 1978. Myasthenic antibodies cross-link acetylcholine receptors to accelerate degradation. N Engl J Med 298: 1116–1122.

30. Sterz, R., R. Hohlfeld, K. Rajki, M. Kaul, K. Heininger, K. Peper, and K. V. Toyka. 1986. Effector mechanisms in myasthenia gravis: end-plate function after passive transfer of IgG, Fab, and F(ab’)2 hybrid molecules. Muscle Nerve 9: 306–312.

31. Howard, F. M., Jr., V. A. Lennon, J. Finley, J. Matsumoto, and L. R. Elveback. 1987. Clinical correlations of antibodies that bind, block, or modulate human acetylcholine receptors in myasthenia gravis. Ann N Y Acad Sci 505: 526–538.

32. Koneczny, I., J. Cossins, P. Waters, D. Beeson, and A. Vincent. 2013. MuSK myasthenia gravis IgG4 disrupts the interaction of LRP4 with MuSK but both IgG4 and IgG1-3 can disperse preformed agrin-independent AChR clusters. PLoS One 8: e80695.

33. Takata, K., P. Stathopoulos, M. Cao, M. Mane-Damas, M. L. Fichtner, E. S. Benotti, L. Jacobson, P. Waters, S. R. Irani, P. Martinez-Martinez, D. Beeson, M. Losen, A. Vincent, R. J. Nowak, and K. C. O’Connor. 2019. Characterization of pathogenic monoclonal autoantibodies derived from muscle-specific kinase myasthenia gravis patients. JCI Insight 4.

34. Huijbers, M. G., D. L. Vergoossen, Y. E. Fillie-Grijpma, I. E. van Es, M. T. Koning, L. M. Slot, H. Veelken, J. J. Plomp, S. M. van der Maarel, and J. J. Verschuuren. 2019. MuSK myasthenia gravis monoclonal antibodies: Valency dictates pathogenicity. Neurol Neuroimmunol Neuroinflamm 6: e547.

35. Stathopoulos, P., A. Kumar, R. J. Nowak, and K. C. O’Connor. 2017. Autoantibody-producing plasmablasts after B cell depletion identified in muscle-specific kinase myasthenia gravis. JCI Insight 2.

36. Fichtner, M. L., C. Vieni, R. L. Redler, L. Kolich, R. Jiang, K. Takata, P. Stathopoulos, P. A. Suarez, R. J. Nowak, S. J. Burden, D. C. Ekiert, and K. C. O’Connor. 2020. Affinity maturation is required for pathogenic monovalent IgG4 autoantibody development in myasthenia gravis. J Exp Med 217.

37. Graus, Y. F., M. H. de Baets, P. W. Parren, S. Berrih-Aknin, J. Wokke, P. J. van Breda Vriesman, and D. R. Burton. 1997. Human anti-nicotinic acetylcholine receptor recombinant Fab fragments isolated from thymus-derived phage display libraries from myasthenia gravis patients reflect predominant specificities in serum and block the action of pathogenic serum antibodies. J Immunol 158: 1919–1929.

38. Cron, M. A., S. Maillard, J. Villegas, F. Truffault, M. Sudres, N. Dragin, S. Berrih-Aknin, and R. Le Panse. 2018. Thymus involvement in early-onset myasthenia gravis. Ann N Y Acad Sci 1412: 137–145.

39. Weis, C. A., B. Schalke, P. Strobel, and A. Marx. 2018. Challenging the current model of early-onset myasthenia gravis pathogenesis in the light of the MGTX trial and histological heterogeneity of thymectomy specimens. Ann N Y Acad Sci 1413: 82–91.

40. Marx, A., F. Pfister, B. Schalke, G. Saruhan-Direskeneli, A. Melms, and P. Strobel. 2013. The different roles of the thymus in the pathogenesis of the various myasthenia gravis subtypes. Autoimmun Rev 12: 875–884.

41. Jiang, R., K. B. Hoehn, C. S. Lee, M. C. Pham, R. J. Homer, F. C. Detterbeck, I. Aban, L. Jacobson, A. Vincent, R. J. Nowak, H. J. Kaminski, S. H. Kleinstein, and K. C. O’Connor. 2020. Thymus-derived B cell clones persist in the circulation after thymectomy in myasthenia gravis. Proc Natl Acad Sci U S A 117: 30649–30660.

42. R. Vander Heiden, J. A., P. Stathopoulos, J. Q. Zhou, L. Chen, T. J. Gilbert, C. R. Bolen, R. J. Barohn, M. M. Dimachkie, E. Ciafaloni, T. J. Broering, F. Vigneault, R. J. Nowak, R. H. Kleinstein, and K. C. O’Connor. 2017. Dysregulation of B Cell Repertoire Formation in Myasthenia Gravis Patients Revealed through Deep Sequencing. J Immunol 198: 1460–1473.

43. Lee, J. Y., P. Stathopoulos, S. Gupta, J. M. Bannock, R. J. Barohn, E. Cotzomi, M. M. Dimachkie, L. Jacobson, C. S. Lee, H. Morbach, L. Querol, J. L. Shan, J. A. Vander Heiden, P. Waters, A. Vincent, R. J. Nowak, and K. C. O’Connor. 2016. Compromised fidelity of B-cell tolerance checkpoints in AChR and MuSK myasthenia gravis. Ann Clin Transl Neurol 3: 443–454.

44. Chen, J., Q. Zheng, C. M. Hammers, C. T. Ellebrecht, E. M. Mukherjee, H. Y. Tang, C. Lin, H. Yuan, M. Pan, J. Langenhan, L. Komorowski, D. L. Siegel, A. S. Payne, and J. R. Stanley. 2017. Proteomic Analysis of Pemphigus Autoantibodies Indicates a Larger, More Diverse, and More Dynamic Repertoire than Determined by B Cell Genetics. Cell Rep 18: 237–247.

45. Sabouri, Z., P. Schofield, K. Horikawa, E. Spierings, D. Kipling, K. L. Randall, D. Langley, B. Roome, R. Vazquez-Lombardi, R. Rouet, J. Hermes, T. D. Chan, R. Brink, D. K. Dunn-Walters, D. Christ, and C. C. Goodnow. 2014. Redemption of autoantibodies on anergic B cells by variable-region glycosylation and mutation away from self-reactivity. Proc Natl Acad Sci U S A 111: E2567–2575.

46. Koers, J., N. I. L. Derksen, P. Ooijevaar-de Heer, B. Nota, F. S. van de Bovenkamp, G. Vidarsson, and T. Rispens. 2019. Biased N-Glycosylation Site Distribution and Acquisition across the Antibody V Region during B Cell Maturation. J Immunol 202: 2220–2228.

47. Anil, R., A. Kumar, S. Alaparthi, A. Sharma, J. L. Nye, B. Roy, K. C. O’Connor, and R. J. Nowak. 2020. Exploring outcomes and characteristics of myasthenia gravis: Rationale, aims and design of registry - The EXPLORE-MG registry. J Neurol Sci 414: 116830.

48. Jiang, R., M. L. Fichtner, K. B. Hoehn, M. C. Pham, P. Stathopoulos, R. J. Nowak, S. H. Kleinstein, and K. C. O’Connor. 2020. Single-cell repertoire tracing identifies rituximab-resistant B cells during myasthenia gravis relapses. JCI Insight 5.

49. Gupta, N. T., K. D. Adams, A. W. Briggs, S. C. Timberlake, F. Vigneault, and S. H. Kleinstein. 2017. Hierarchical Clustering Can Identify B Cell Clones with High Confidence in Ig Repertoire Sequencing Data. J Immunol 198: 2489–2499.

50. Ye, J., N. Ma, T. L. Madden, and J. M. Ostell. 2013. IgBLAST: an immunoglobulin variable domain sequence analysis tool. Nucleic Acids Res 41: W34–40.

51. Gupta, N. T., J. A. Vander Heiden, M. Uduman, D. Gadala-Maria, G. Yaari, and S. H. Kleinstein. 2015. Change-O: a toolkit for analyzing large-scale B cell immunoglobulin repertoire sequencing data. Bioinformatics 31: 3356–3358.

52. Yaari, G., J. A. Vander Heiden, M. Uduman, D. Gadala-Maria, N. Gupta, J. N. Stern, K. C. O’Connor, D. A. Hafler, U. Laserson, F. Vigneault, and S. H. Kleinstein. 2013. Models of somatic hypermutation targeting and substitution based on synonymous mutations from high-throughput immunoglobulin sequencing data. Front Immunol 4: 358.

53. Hoehn, K. B., J. A. Vander Heiden, J. Q. Zhou, G. Lunter, O. G. Pybus, and S. H. Kleinstein. 2019. Repertoire-wide phylogenetic models of B cell molecular evolution reveal evolutionary signatures of aging and vaccination. Proc Natl Acad Sci U S A 116: 22664–22672.

54. Yu, G., T. T. Lam, H. Zhu, and Y. Guan. 2018. Two Methods for Mapping and Visualizing Associated Data on Phylogeny Using Ggtree. Mol Biol Evol 35: 3041–3043.

55. Srzentic, K., L. Fornelli, Y. O. Tsybin, J. A. Loo, H. Seckler, J. N. Agar, L. C. Anderson, D. L. Bai, A. Beck, J. S. Brodbelt, Y. E. M. van der Burgt, J. Chamot-Rooke, S. Chatterjee, Y. Chen, D. J. Clarke, P. O. Danis, J. K. Diedrich, R. A. D’Ippolito, M. Dupre, N. Gasilova, Y. Ge, Y. A. Goo, D. R. Goodlett, S. Greer, K. F. Haselmann, L. He, C. L. Hendrickson, J. D. Hinkle, M. V. Holt, S. Hughes, D. F. Hunt, N. L. Kelleher, A. N. Kozhinov, Z. Lin, C. Malosse, A. G. Marshall, L. Menin, R. J. Millikin, K. O. Nagornov, S. Nicolardi, L. Pasa-Tolic, S. Pengelley, N. R. Quebbemann, A. Resemann, W. Sandoval, R. Sarin, N. D. Schmitt, J. Shabanowitz, J. B. Shaw, M. R. Shortreed, L. M. Smith, F. Sobott, D. Suckau, T. Toby, C. R. Weisbrod, N. C. Wildburger, J. R. Yates, 3rd, S. H. Yoon, N. L. Young, and M. Zhou. 2020. Interlaboratory Study for Characterizing Monoclonal Antibodies by Top-Down and Middle-Down Mass Spectrometry. J Am Soc Mass Spectrom 31: 1783–1802.

56. Fornelli, L., K. Srzentic, R. Huguet, C. Mullen, S. Sharma, V. Zabrouskov, R. T. Fellers, K. R. Durbin, P. D. Compton, and N. L. Kelleher. 2018. Accurate Sequence Analysis of a Monoclonal Antibody by Top-Down and Middle-Down Orbitrap Mass Spectrometry Applying Multiple Ion Activation Techniques. Anal Chem 90: 8421–8429.

57. Stathopoulos, P., A. Chastre, P. Waters, S. Irani, M. L. Fichtner, E. S. Benotti, J. M. Guthridge, J. Seifert, R. J. Nowak, J. H. Buckner, V. M. Holers, J. A. James, D. A. Hafler, and K. C. O’Connor. 2019. Autoantibodies against Neurologic Antigens in Nonneurologic Autoimmunity. J Immunol 202: 2210–2219.

